# Complement C3 Deficiency Enhances Renal Leptospiral Load and Inflammation While Impairing T Cell Differentiation During Chronic *Leptospira interrogans* Infection

**DOI:** 10.1101/2025.03.31.646275

**Authors:** Leonardo Moura Midon, Amaro Nunes Duarte Neto, Ana Maria Gonçalves da Silva, Marcos Bryan Heinemann, Suman Kundu, Maria Gomes-Solecki, Lourdes Isaac

**Affiliations:** Department of Immunology, Institute of Biomedical Sciences, University of São Paulo, São Paulo, Brazil; Department of Pathology, Faculty of Medicine, University of São Paulo, São Paulo, Brazil; Tropical Medicine Institute, Faculty of Medicine, University of São Paulo, São Paulo, Brazil; Bacterial Zoonosis Laboratory, Medicine Veterinary School, University of São Paulo; Department of Microbiology, Immunology and Biochemistry, University of Tennessee Health Science Center, Memphis, USA

**Keywords:** Complement System, leptospirosis, C3, chronic kidney fibrosis, pathology, T cell maturation

## Abstract

Leptospirosis is a neglected zoonotic disease caused by pathogenic *Leptospira* spp., affecting an estimated of one million people annually and resulting in approximately 60,000 fatalities. The disease can lead to hepatic, renal, and pulmonary dysfunctions and may contribute to the development of chronic kidney disease.

The Complement System plays a crucial role in eliminating bacteria by generating opsonins, anaphylotoxins, that degranulate mastocytes and basophils, and attract immune cells to the infection site, among other important functions.

This study aimed to investigate the role of C3, the central protein of the Complement System, *in vivo* during chronic infection of *L interrogans* serovar Copenhageni strain FIOCRUZ L1-130 (LIC). C57BL/6 wild-type (WT) and C3 knockout (C3KO) mice were infected with 10^8^ LIC and monitored at 15, 30, 60, 90 and 180 days’ post-infection (d.p.i.).

LIC-infected C3KO mice exhibited a significantly higher leptospiral load in the kidneys compared to WT counterparts. Both groups showed local inflammation at 15 and 30 d.p.i., but only C3KO LIC-infected mice had a higher number of *Leptospira* DNA copies at 30 d.p.i. At the same time point, C3KO LIC-infected mice developed a larger fibrotic area than WT mice. Interestingly, independent of C3, mice pre-treated with a nephrotoxic drug increased the renal inflammatory response; however, this pretreatment did not affect the local leptospiral load in infected mice. Additionally, levels of specific IgG2b and IgG3 antibodies were significantly higher in LIC-infected C3KO mice compared to WT mice.

Proteomic analysis showed lower levels of C5/C5a in WT and C3KO LIC-infected mice, as well as in C3KO control mice. M-CSF and SDF-1 cytokine levels were reduced in both LIC-infected groups. Furthermore, naïve T lymphocytes count (both CD4^+^ and CD8^+^) were higher in LIC-infected C3KO mice, whereas effector CD8^+^ T lymphocyte numbers declined during LIC infection - a phenomenon more pronounced in C3KO mice.

Overall, this study demonstrates that during LIC infection, the absence of C3 does not affect mouse survival but leads to increased renal leptospiral load and fibrosis. Additionally, it highlights the crucial role of C3 in supporting the maturation and differentiation of T lymphocytes into pre-effector cells, underscoring its importance as a key link between the innate and adaptive immune responses in leptospirosis.

**Author Summary:** Leptospirosis is an infectious disease with approximately one million new cases annually and is responsible for about 5% of deaths, particularly in underdeveloped countries. Our objective is to understand the immune response in leptospirosis, specifically the role of the Complement C3 protein. In this study, we observed that C3 deficiency is associated with a higher renal leptospiral load and an increased incidence of renal fibrosis after one month of chronic infection. Notably, C3 also influences the differentiation of helper and cytotoxic T lymphocytes into effector cells, potentially contributing to the increased severity of chronic leptospirosis observed in C3-deficient animals.

## Introduction

Leptospirosis is a neglected disease and one of the most important zoonoses worldwide, affecting an estimated one million people and causing 60,000 deaths annually [1]. It is caused by the pathogenic *Leptospira* spp. [2, 3] occurs mainly in developing countries with tropical and mild climates with poor sanitary conditions. In urban centers, infected rodents, such as *Rattus novergicus* (brown rat), serve as primary reservoirs of *Leptospira*, shedding bacteria through their urine [4, 5].

Humans are accidental hosts who become infected upon contact with contaminated water or soil. *Leptospira* can penetrate the host through mucosa or damaged skin. Most cases are asymptomatic or present mild symptoms such as fever, headache, muscle pain, nausea, and vomiting during the blood dissemination phase (the acute phase). These symptoms overlap those of other acute infections, leading to underreporting of leptospirosis [3, 4].

Leptospirosis can lead to multiple organ dysfunction, particularly affecting the liver, kidneys, and lungs [6]. In severe cases, such as Weil’s syndrome, complications may include jaundice, kidney and liver failure, internal hemorrhage, and pulmonary distress, which can be fatal [6, 7]. Pulmonary hemorrhage associated with leptospirosis is another possible severe manifestation [8], with a high mortality rate (30-60%) and the potential to cause fulminant death in a short period [9, 10]. Leptospirosis may also be linked to chronic kidney disease of unknown etiology in endemic areas of Southeast Asia and Latin America [11].

C3 is the most abundant Complement System protein [12] in the serum and plays a key role in activating the Classical, Lectin and Alternative Pathways, which converge at the Terminal Pathway to generate the membrane attack complex (MAC) on pathogen surfaces, potentially causing osmotic imbalance and destruction. C3 mediates various immune functions, including: *i)* attracting immune cells to the site of inflammation site via its C3a fragment; *ii)* opsonizing pathogens through C3b and iC3b fragments, facilitating phagocytosis via Complement Receptors (CR); *iii)* activating mast cells and basophils, leading to release of inflammatory mediators; and *iv)* supporting B-lymphocyte activation and antibody production via the C3d fragment [reviewed in 13; 14].

C3 deficiency is associated with susceptibility to severe and recurrent bacterial infections [14, 15], as well as chronic renal diseases, such as membranoproliferative glomerulonephritis [16]. Dysregulated activation of C3, however, can lead to tissue damage, particularly in the kidney [17–19].

Pathogenic *Leptospira* spp. have evolved immune evasion mechanisms to circumvent the Complement System [20] and disseminate in the host by: *i)* binding to host Complement regulatory proteins [21]; *ii)* acquiring host proteases capable of cleaving C3 and C5 [22]; and, secreting metalloproteases such as thermolysin [23, 24] and leptolysin [25] that cleave Complement proteins and macrophage surface molecules necessary to phagocytosis [26].

In a previous study conducted by our group [27], we investigated the role of C3 during the acute phase of infection with *L. interrogans* serovar Kennewicki type Pomona Fromm in C57BL/6 mice wild-type (WT) and C3-deficient (C3KO) mice. The absence of C3 was associated with significantly higher leptospiral loads in the kidney, liver, spleen, and urine compared to WT mice at 3 and 6 days post-infection (d.p.i.). Interstitial nephritis was observed in C3KO mice at 15 d.p.i.

In this study, we investigated the role of C3 in a chronic model of leptospirosis using *L. interrogans* serovar L1-130 FIOCRUZ at multiple time points, assessing its importance in kidney fibrosis formation, an immune response against this pathogen.

## Results

### Survival and *Leptospira* renal colonization in infected mice

WT and C3KO mice were infected (i/p) with 10^8^ LIC or PBS (control) and monitored at 15, 30, 60, 90, and 180 d.p.i. All groups of mice survived, and body weight differences were not significant when comparing LIC-infected WT and C3KO mice even at 180 d.p.i. Splenomegaly was observed in infected mice at multiple time points when compared to uninfected controls. However, in the absence of C3, this increase was significantly attenuated at 30 d.p.i. compared to infected WT mice (**S1 Fig**). The presence of LIC in kidney tissue was confirmed by immunohistochemistry analysis (**S2 Fig**) and quantified by qPCR (**Fig 1**). A higher leptospiral load was observed in the kidneys of LIC-infected C3KO mice at 30 d.p.i compared to WT counterparts.

**Fig 1.**
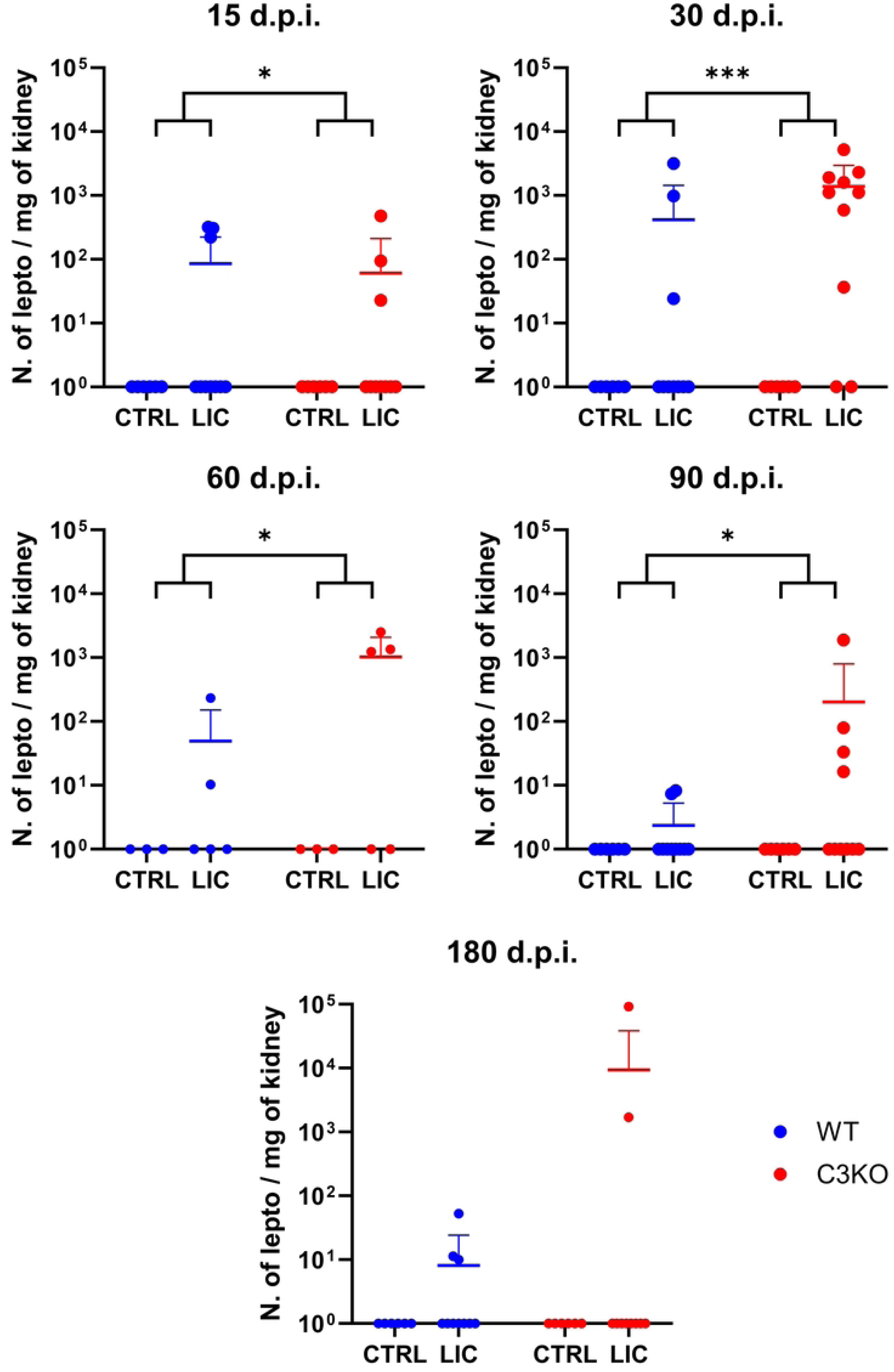
Quantification of kidney leptospiral load. Genomic DNA was extracted from the kidneys of WT and C3KO mice inoculated with PBS (control; CTRL) or 10^8^ *L interrogans* serovar Copenhageni strain FIOCRUZ L1-130 (LIC) (i/p) and monitored for 15, 30, 60, 90, and 180 days post-infection (d.p.i.). Leptospiral load was quantified using a standard qPCR curve and expressed as the number of leptospires (N.of lepto) per mg of kidney tissue. Statistical analysis was performed using Kruskal-Wallis nonparametric test, followed by Dunn’s post-hoc test. 15 d.p.i: CTRL – LIC *p* = 0.040; 30 d.p.i: CTRL – LIC *p* = 0.002 and WT – C3KO *p* = 0.007; 60 d.p.i.: CTRL – LIC *p* = 0.025; 90 d.p.i.: CTRL – LIC *p* = 0.034; 180 d.p.i.: CTRL – LIC *p* = 0.062. Mice were obtained from the Animal Care Unit from ICB-USP. CTRL group (n=3) and LIC-infected group (n+5). Infections at 15, 30, 90 and 180 d.p.i. were repeated twice, while infection at 60 d.p.i. was performed once.

### Renal injury during *Leptospira* infection

Histopathological analysis of kidney samples (**Fig 2**) showed that non-infected WT and C3KO mice were apparently healthy, with no inflammatory cell infiltration. In contrast, kidneys from both groups of LIC-infected mice at 15 d.p.i. and 30 d.p.i exhibited inflammatory cell infiltrates near the vascular endothelium, fibrosis, and nephritis. Although no significant differences were observed between LIC-infected WT and C3KO mice, C3KO mice displayed persistent kidney injury even after 180 d.p.i. when compared to non-infected C3KO control (**Fig 3**). On the other hand, LIC-infected WT mice showed no significant tissue damage after two, three or six months of infection compared to non-infected WT mice.

**Fig 2.**
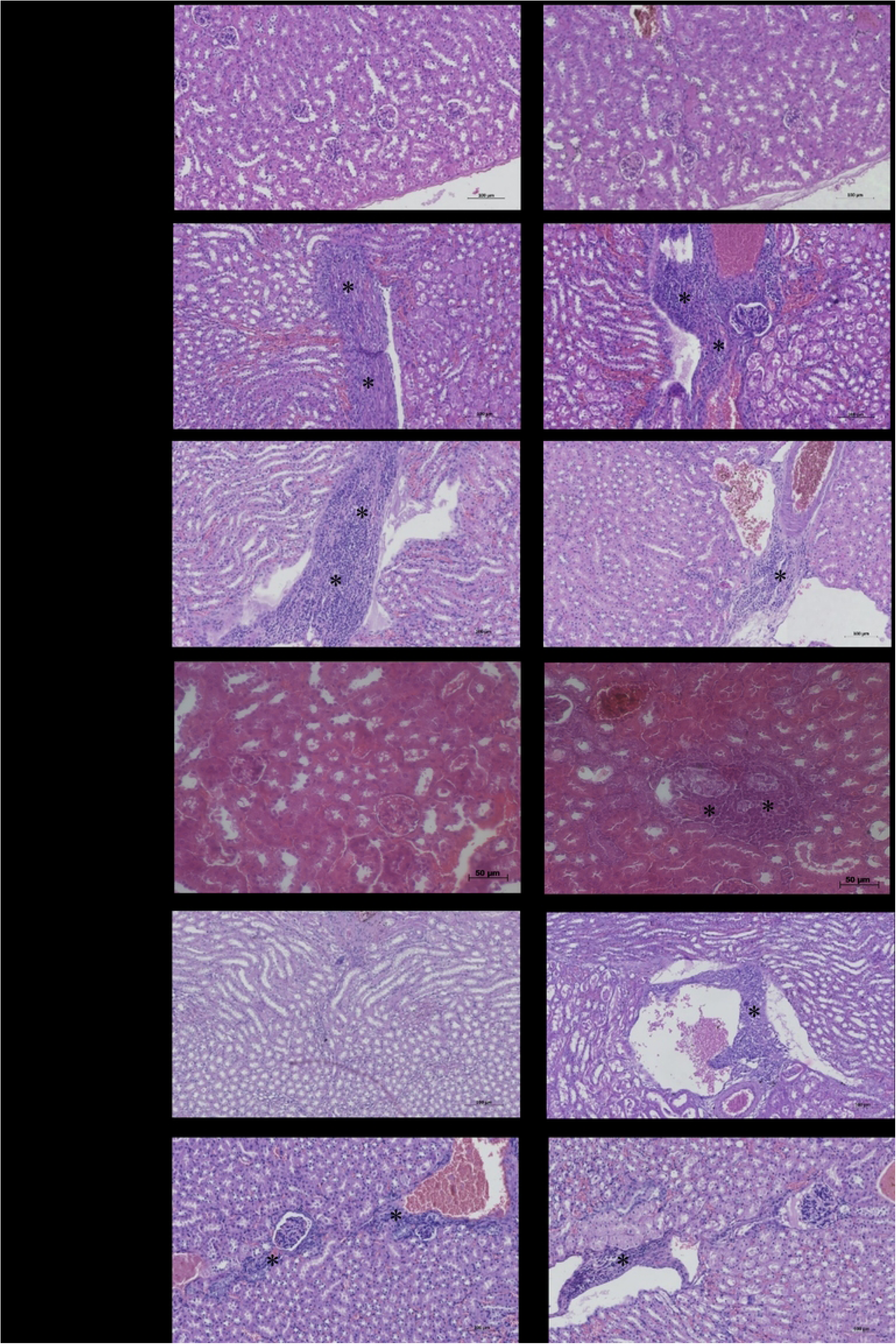
Kidney histopathological analysis after infection with pathogenic leptospires. WT and C3KO mice were inoculated with PBS (control; CTRL) or 10^8^ *L interrogans* serovar Copenhageni strain FIOCRUZ L1-130 (LIC) (i/p) and monitored for 15, 30, 60, 90, and 180 days post-infection (d.p.i.). Kidney sections were stained with hematoxylin-eosin (HE). Inflammatory infiltrates (*), nephritis, and fibrosis were observed in both groups of LIC-infected mice. Images are shown at 200x magnification. Mice were obtained from the Animal Care Unit from ICB-USP.

**Fig 3.**
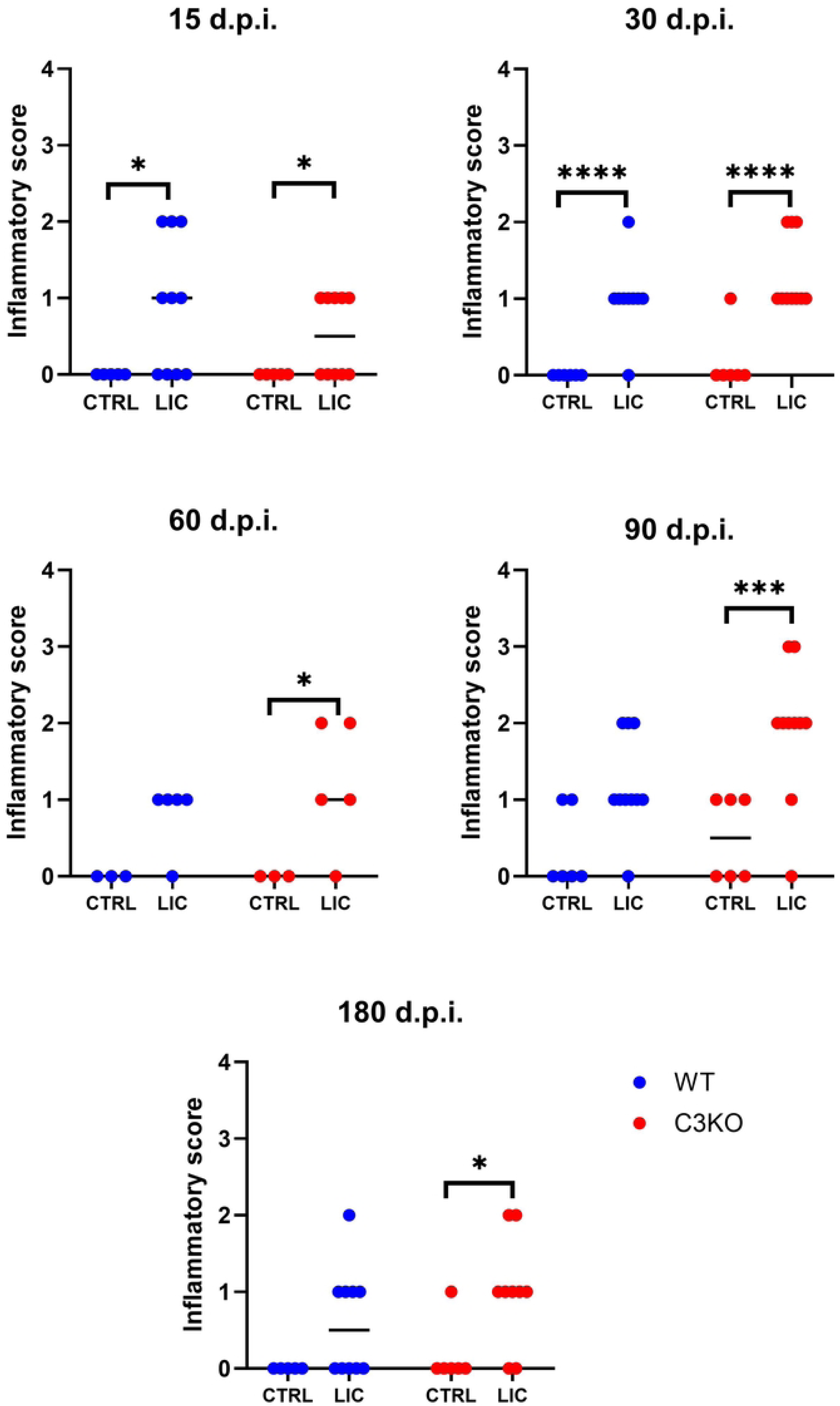
Renal inflammatory score. WT or C3KO mice were inoculated with PBS (control; CTRL) or 10^8^ *L interrogans* serovar Copenhageni strain FIOCRUZ L1-130 (LIC) (i/p) and monitored for 15, 30, 60, 90, and 180 days post-infection (d.p.i.). The score was classified as follows: 0 – no alterations; 1 - < 25% of fibrosis, nephritis, and inflammatory infiltrates; 2 – 25% < and < 50% of fibrosis, nephritis and inflammatory infiltrates; and 3 - 50% < % of fibrosis, nephritis, and inflammatory infiltrates. Statistical analysis was performed using Kruskal-Wallis test (as the values are ordinal numbers), followed by Dunn’s post-hoc test. \**p* < 0.05; \*\*\**p* < 0.001, and \*\*\*\**p* < 0.0001. Mice were obtained from the Animal Care Unit from ICB-USP. Mice were obtained from the Animal Care Unit from ICB-USP. CTRL group (n=3) and LIC-infected group (n+5). Infections at 15, 30, 90 and 180 d.p.i. were repeated twice, while infection at 60 d.p.i. was performed once.

### Parameters of renal function and fibrosis formation during *Leptospira* infection

Renal function was assessed by measuring serum levels of urea, uric acid, and creatinine levels (**Fig 4**) in both infected and non-infected WT and C3KO mice. The lack of C3 influenced uric acid levels leading to a reduction in serum levels at 15 and 30 d.p.i., and subsequently resulted in a decrease in serum urea levels at 180 d.p.i., with significant differences observed when compared to WT-infected mice. In contrast, serum creatinine levels remained similar between WT and C3KO-infected animals.

**Fig 4.**
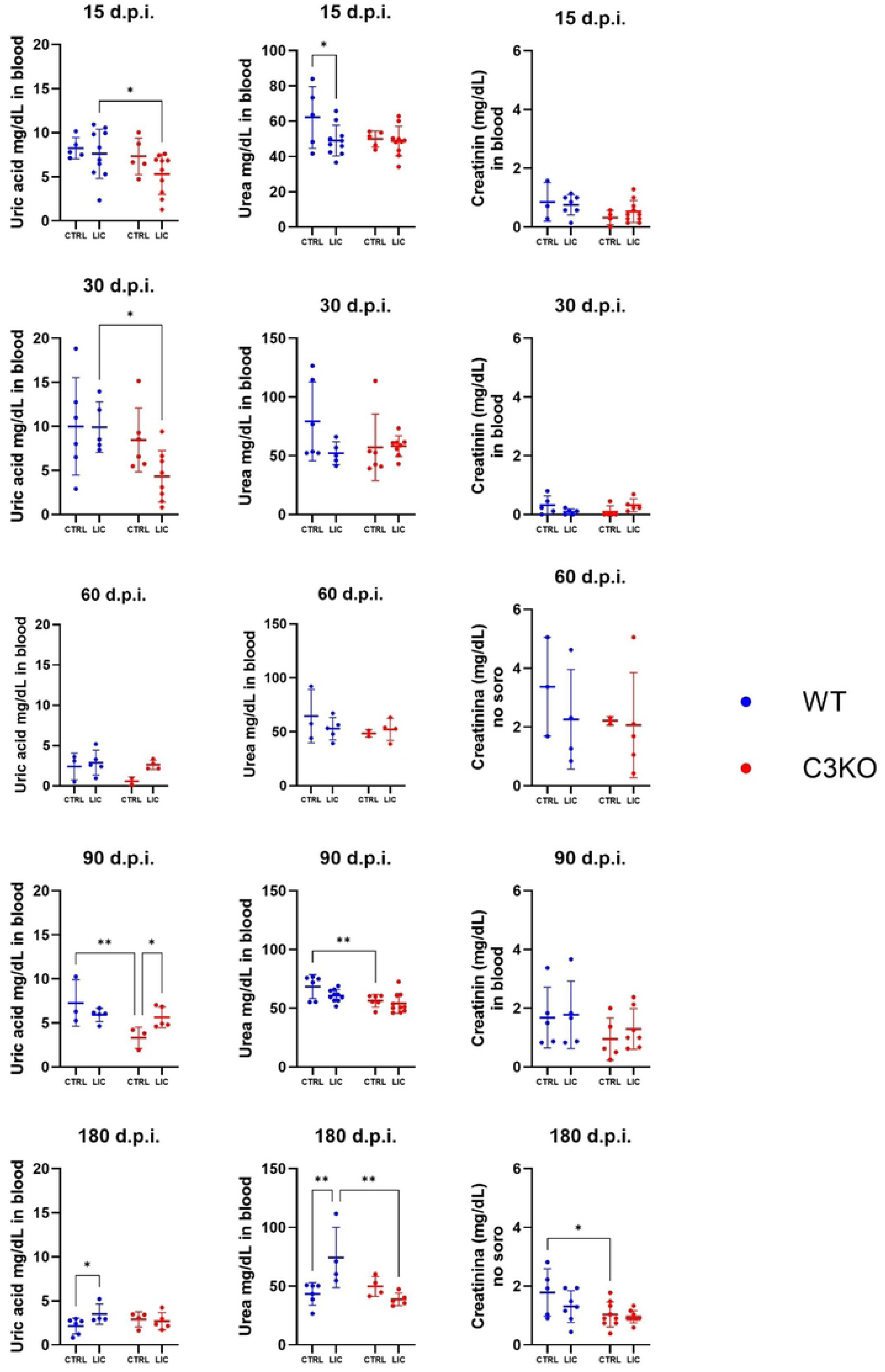
Serum levels of uric acid, urea, and creatinine. WT or C3KO mice were inoculated with PBS (control; CTRL) or with of 10^8^ *L interrogans* serovar Copenhageni strain FIOCRUZ L1-130 (LIC) (i/p). Data are represented as mean ± S.D. Each dot represents one animal. Statistical analysis was performed using two-way ANOVA, followed by Tukey’s test, with familiar α of 0.95. \**p*< 0.05; and \*\**p*< 0.01. Mice were obtained from the Animal Care Unit from ICB-USP. CTRL group (n=3) and LIC-infected group (n+5). Infections at 15, 30, 90 and 180 days post infection (d.p.i.) were repeated twice, while infection at 60 d.p.i. was performed once.

Additionally, fibrosis formation in LIC-infected mice was monitored. The absence of C3 was significantly associated with larger areas of kidney fibrosis at 30 d.p.i. compared to WT mice (**Fig 5**). Interestingly, this result correlates with the highest leptospiral load observed at 30 and 60 d.p.i. in LIC-infected C3KO mice, gradually decreasing thereafter (**Fig 1**).

**Fig 5.**
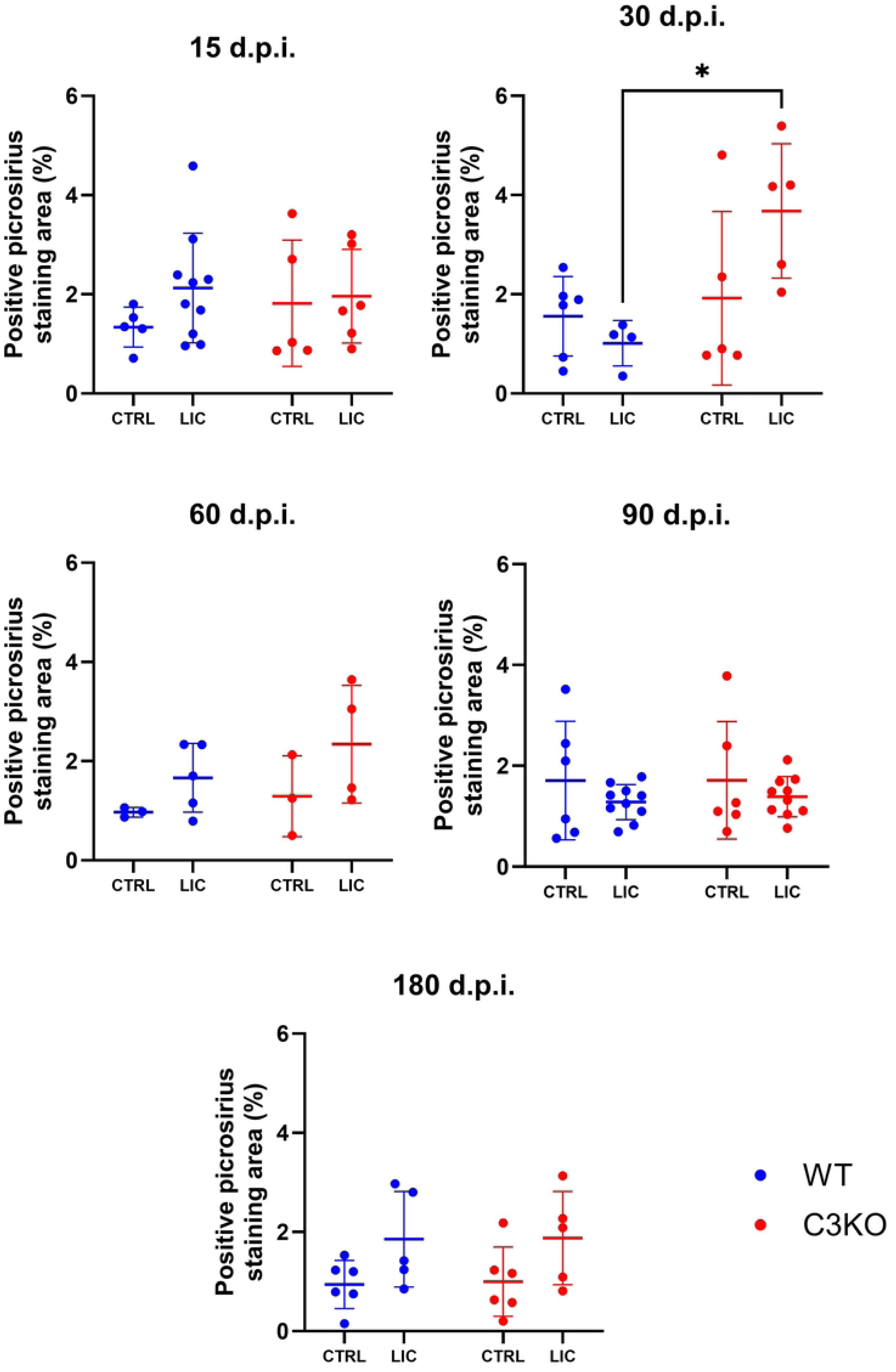
Fibrosis area percentage during LIC infection. WT or C3KO mice were FIOCRUZ L1-130 (LIC) (i/p) and followed from 15 days post-infection (d.p.i.) up to six months (180 d.p.i.). Sirius Red-stained kidney sections were scanned with the ZEISS Axioscan 7 slide scanner. Red intensity was quantified using Image J. Statistical analysis was performed using two-way ANOVA, followed by Tukey’s test. \**p*< 0.05. Mice were obtained from the Animal Care Unit from ICB-USP. CTRL group (n=3) and LIC-infected group (n+5). Infections at 15, 30, 90 and 180 d.p.i. were repeated twice, while infection at 60 d.p.i. was performed once.

We also analyzed the mRNA production of fibrosis-related genes, including fibronectin (*FN1*), collagen-1 (*COL1A1*) and alpha-smooth muscle actin (*α-SMA*). However, neither C3 deficiency nor LIC infection appeared to significantly alter these parameters (**S3 Fig**).

### *Leptospira* infection in cisplatin-treated mice

To further investigate whether pre-existing kidney lesions induced by a nephrotoxic agent exacerbate *Leptospira* infection in the absence of C3, we administered cisplatin - a chemotherapeutic drug with high nephrotoxicity - to mice and later assessed its impact in conjunction with LIC infection (**Fig 6**). While C3 was not required to influence cisplatin-induced kidney damage, LIC infection resulted in a significantly higher inflammation score in animals pre-treated with cisplatin. Moreover, a significantly increased leptospiral load was observed in C3KO LIC-infected mice without cisplatin treatment compared to WT LIC-infected mice.

**Fig 6.**
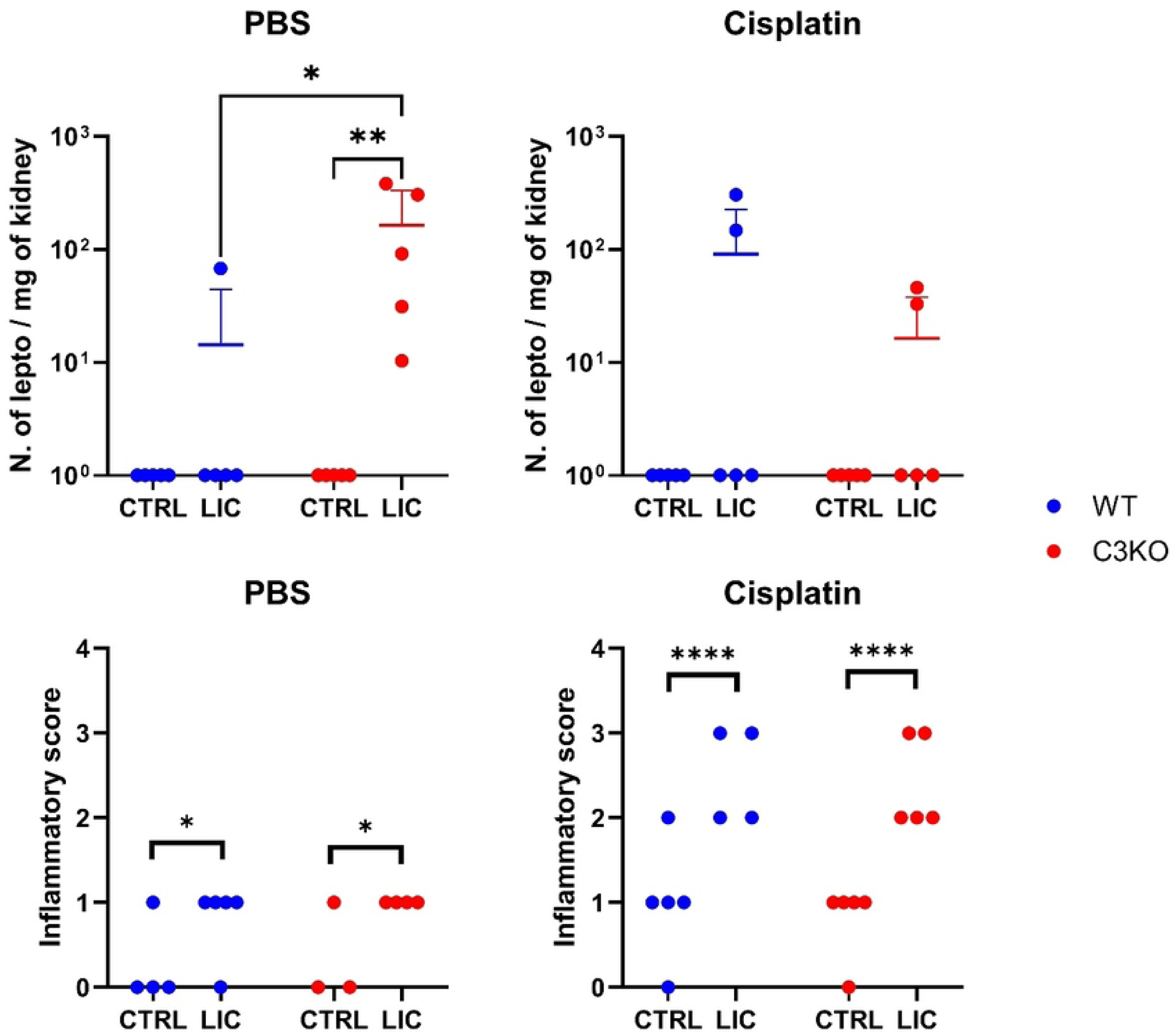
Kidney leptospiral load and inflammatory score after cisplatin treatment. WT and C3KO mice injected with PBS (CTRL) or cisplatin. Thirty days later, the mice were subsequently inoculated with PBS (control; CTRL) or 10^8^ *L interrogans* serovar Copenhageni strain FIOCRUZ L1-130 (LIC) (i/p). Genomic DNA was extracted from the kidneys to check for the leptospiral load, quantified using a standard qPCR curve. The inflammatory score was classified as follows: 0 – no alterations; 1 - < 25% of fibrosis, nephritis, and inflammatory infiltrates; 2 – 25% < and < 50% of fibrosis, nephritis, and inflammatory infiltrates; and 3 - 50% < % of fibrosis, nephritis, and inflammatory infiltrates. Statistical analysis was performed using Kruskal-Wallis. \**p* < 0.05; and \*\*\*\**p* < 0.001. Mice were obtained from the Animal Care Unit from ICB-USP. Mice were obtained from the Animal Care Unit from ICB-USP. Each group n=5.

### Cytokine production

A proteome profile array was used to analyze the presence of several cytokines (including chemokines) and Complement C5/C5a protein in pooled serum (n = 3-4 per group). The array measured cytokines TNF-α, IFN-γ, IL-1α, IL-1β, IL-1ra, IL-2, IL-3, IL-4, IL-5, IL-6, IL-7, IL-10, IL-12 p70, IL-13, IL-16, IL-17, IL-23 and IL-27 (**Fig 7**), including the chemokines BLC, G-CSF, GM-CSF, I-309, Eotaxin, sICAM-1, IP-10, I-TAC, KC, M-CSF, JE, MCP-5, MIG, MIP-1α, MIP-1β, MIP-2, RANTES, SDF-1, TARC, TIMP-1 and TREM-1 (**Fig 7**). At 30 d.p.i., the relative amounts of all cytokines from infected groups (WT or KO) were lower than in their respective controls. Similarly, chemokine levels were generally reduced in infected groups compared to controls, with the exception of IP-10 and KC in WT mice and sICAM-1 in C3KO mice, which remained unchanged. Between uninfected groups (WT *vs* C3KO), the profiles of cytokines TNF-α, IFN-γ, IL-α, IL-1Ra, IL-3, IL-16, IL-17, IL-27 and chemokines BLC, G-CSF, IP-10, KC, M-CSF, JE, MIG, RANTES, SDF-1, TIMP-1 and TREM-1 were higher in the C3KO group. The relative level of C5/C5a was higher in the WT uninfected group compared to the LIC infected counterpart, but this difference was not detected in the C3KO groups.

**Fig 7.**
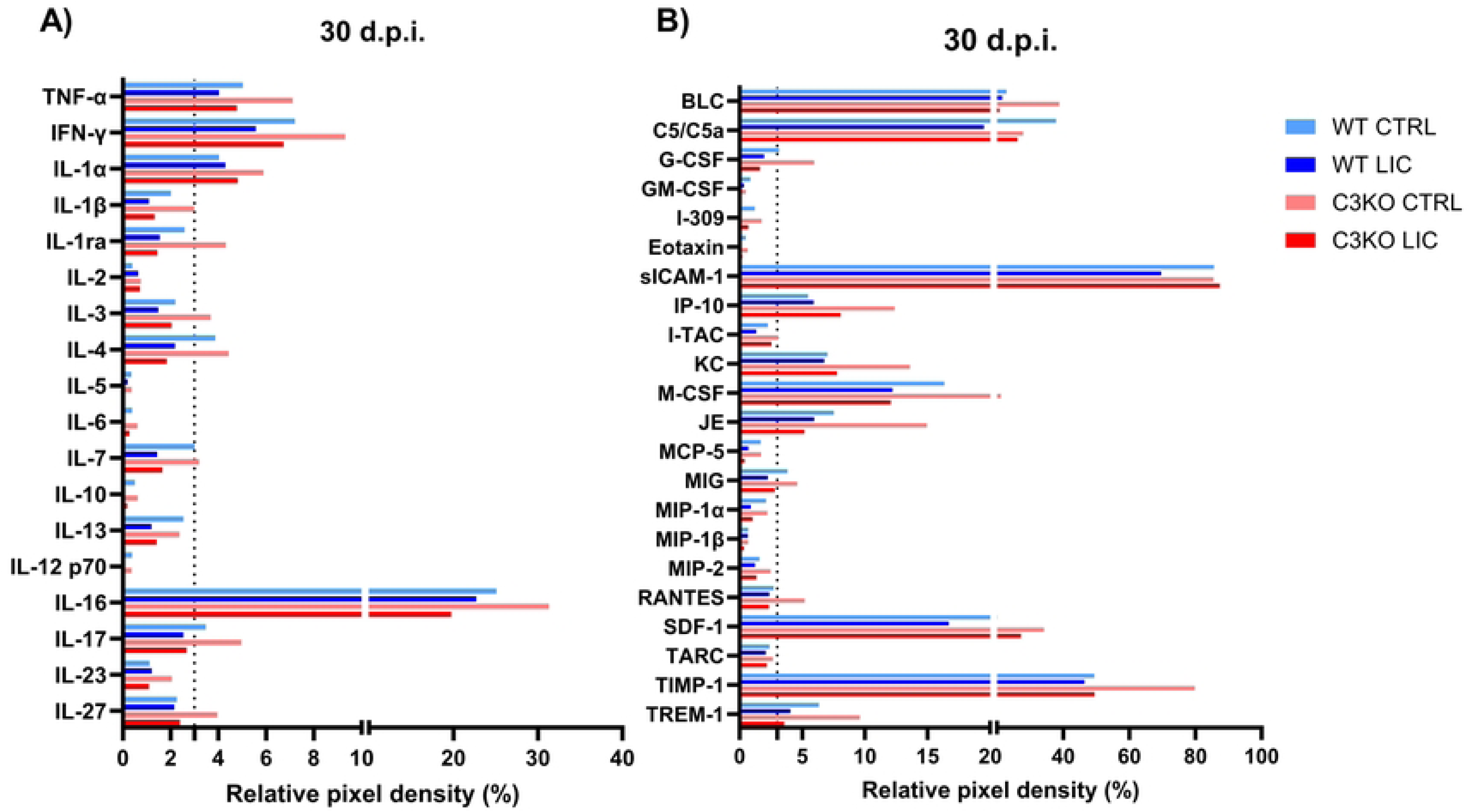
Serum cytokine and acute-phase protein expression in LIC-infected mice at 30 d.p.i.. Chemiluminescence was measured on the pixel density mean of two spots using Quickspot software (ARY006 kit). Values were normalized by subtracting control reference dots (PBS) and calibrating with six positive control dots. Results were considered different if there was a ± 3% difference. Cytokine levels below 3% (dotted line) were considered 0%. Mice were obtained from the Animal Care Unit from UTHSC. CTRL group (n=3) and LIC-infected group (n=5).

### Production of specific anti-LIC antibodies

Both groups of mice were infected with viable LIC, and the production of specific anti-LIC antibodies was assessed in serum at 15 d.p.i.. (**Fig 8**). Both WT and C3KO LIC-infected mice exhibited significantly higher serum levels of IgM (**Fig 8**) compared to non-infected control mice. No significant differences were observed in the serum of specific IgM, total IgG and IgG1 between WT and C3KO LIC-infected mice. However, in the absence of C3, LIC-infected mice produced higher levels of IgG2b and IgG3 compared to WT counterparts, likely due to prolonged leptospiral survival in these deficient animals.

**Fig 8.**
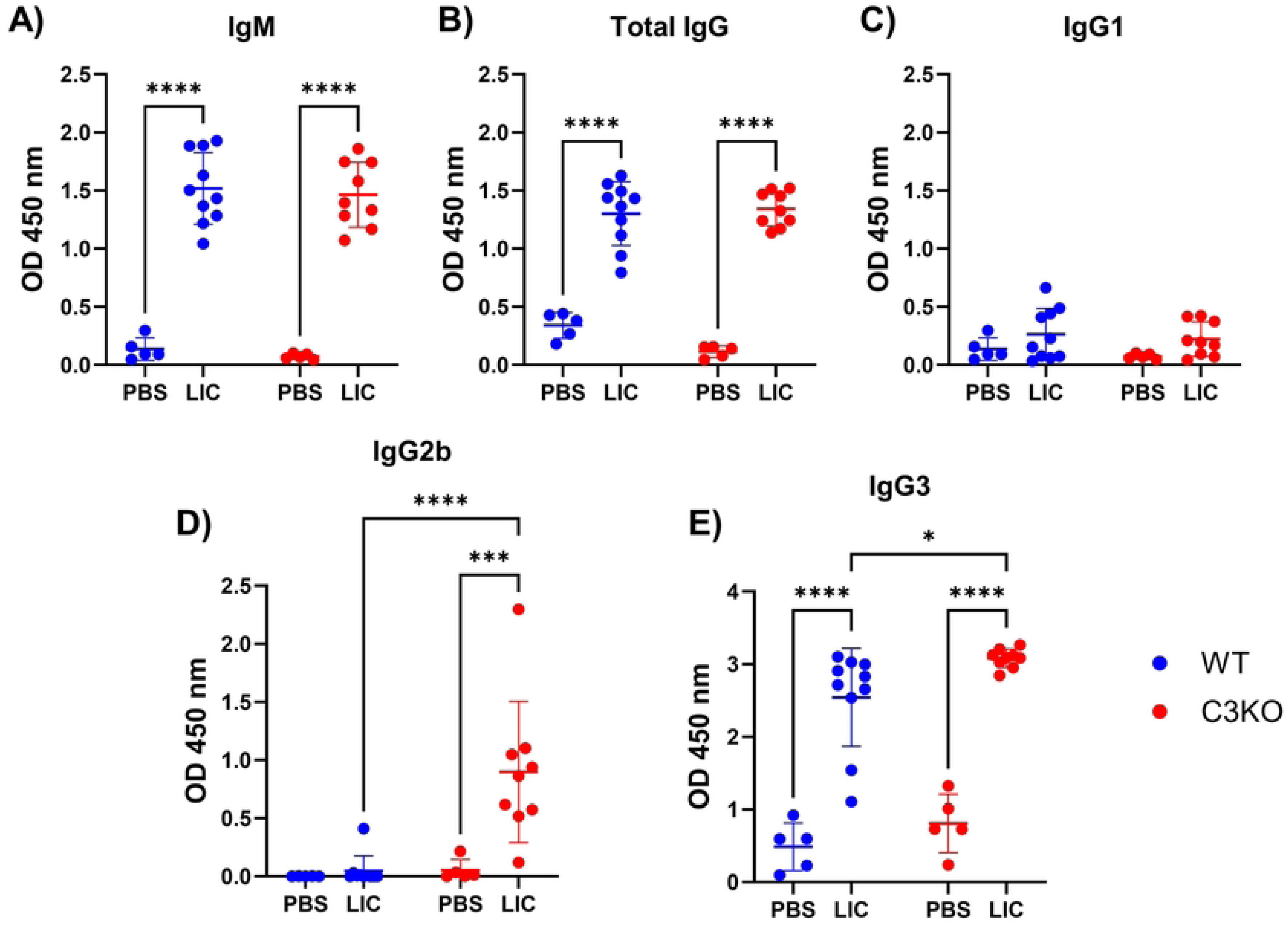
Quantification of specific antibodies against LIC. WT or C3KO mice were inoculated with PBS (control, CTRL) or 10^8^ *L interrogans* serovar Copenhageni strain FIOCRUZ L1-130 (LIC) (i/p). After 15 d.p.i., serum was obtained and levels of specific IgM, total IgG, and other IgG subclasses anti-LIC were quantified by ELISA. Wells were coated with 10^6^ heat-killed LIC and incubated with diluted mice serum (1:100). Secondary antibodies against **(A)** IgM, **(B)** total IgG, **(C)** IgG1, **(D)** IgG2b, and **(e)** IgG3 were used (1:5000). Each dot represents one animal, (n = 5-6 for PBS groups; n = 5-10 for infected groups). Statistical analysis was performed using two-way ANOVA followed by Tukey’s test, with familiar α of 0.95. *p-*values: *< 0.05; \*\*\**p*< 0.001; \*\*\*\**p*< 0.0001. Mice were obtained from the Animal Care Unit from ICB-USP. CTRL group (n=5) and LIC-infected group (n=9-10). Data represent two independent experiments.

### Effect of C3 on effector T cell populations during *Leptospira* infection

Flow cytometry analysis of immune cells in the kidney, lymph nodes, and spleen revealed no significant differences in the percentage of lymphoid cells (CD45^+^), B lymphocytes (CD19^+^) and T (CD3^+^) lymphocytes among the four experimental groups (non-infected *vs* LIC-infected; WT *vs* C3KO) (**Fig 9**, **S4 Fig**).

**Fig 9.**
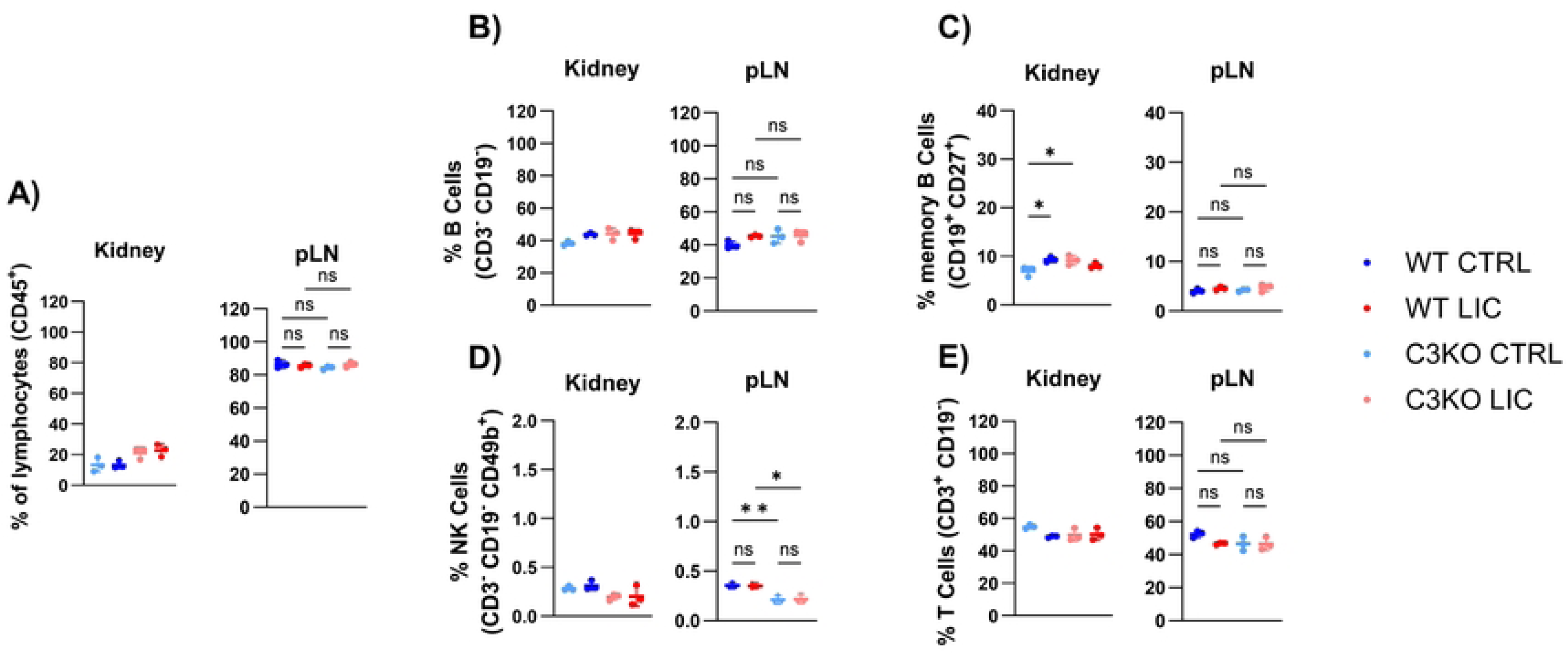
Immune cell populations in the kidney and peripheral lymph nodes (pLN). WT or C3KO mice were inoculated with PBS (control; CTRL) or with 10^8^ *L interrogans* serovar Copenhageni strain FIOCRUZ L1-130 (LIC) (i/p). Graphs represent the percentage of CD45^+^ T and B lymphocytes, their subpopulations, and NK cells after 30 days post-infection. Each dot represents one animal (n =3 per group). Statistical analysis was performed using two-way ANOVA, followed by Tukey’s test, with familiar α of 0.95. *p-*values: *< 0.05; **< 0.01; ns = non-significant. Mice were obtained from the Animal Care Unit from UTHSC.

However, in the absence of C3, a lower proportion of effector T cells was detected in both kidney and lymph node, but not in the spleen, of infected mice, suggesting a delay in the differentiation of naïve T cells into effector T cytotoxic cells. A similar trend was observed in early effector T helper lymphocytes, indicating that C3 may play a crucial role in promoting local T cell differentiation (**Fig 10**). Additionally, NK cells were present in proportionally low in the lymph nodes of C3KO mice; however, this difference was relatively minor compared to WT counterparts (**Fig 9**).

**Fig 10.**
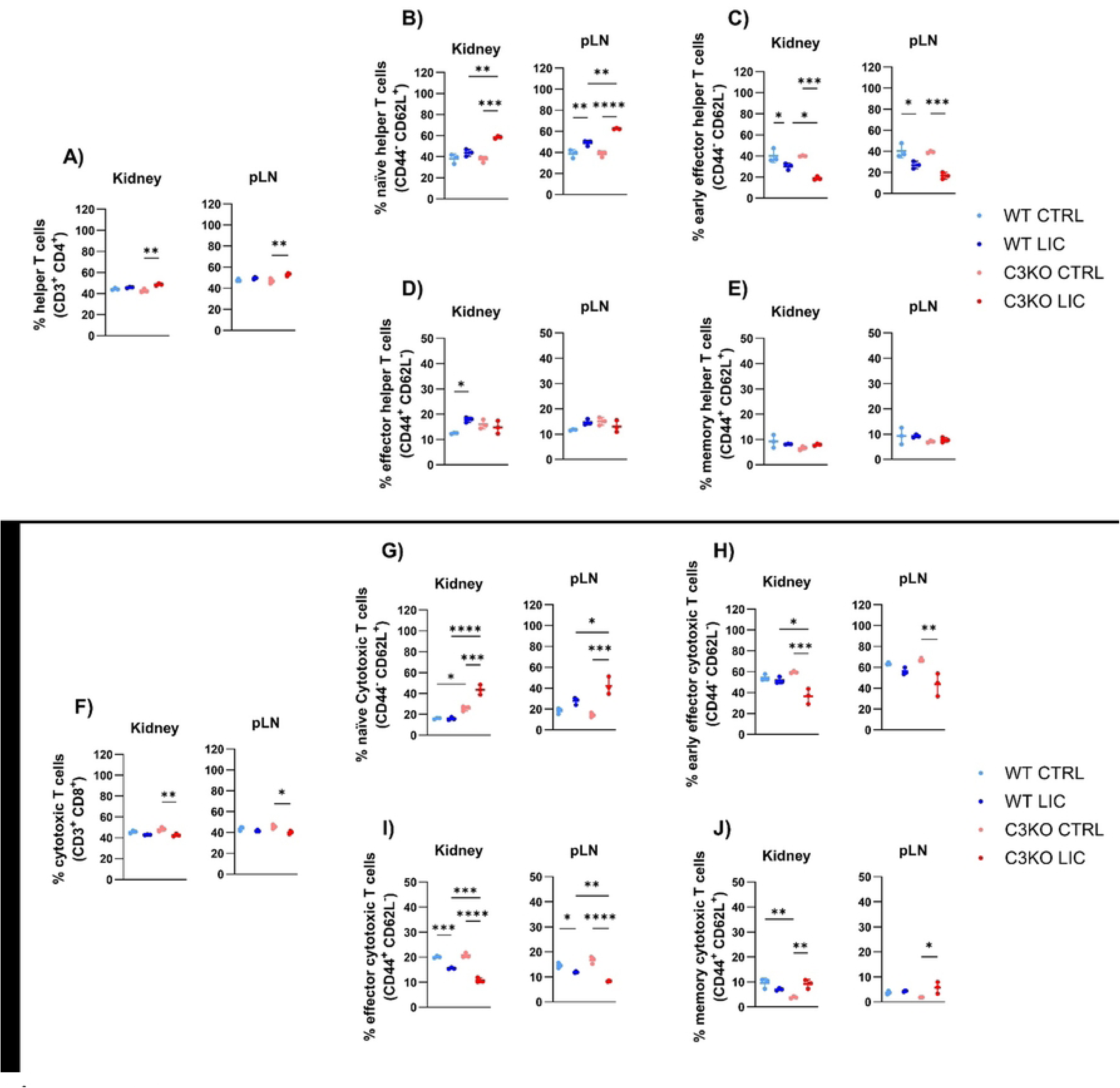
Helper T (CD4^+^) and cytotoxic T (CD8) cell populations in the kidney and peripheral lymph nodes (pLN). WT or C3KO mice were inoculated with PBS (control; CTRL) or with of 10^8^ *L interrogans* serovar Copenhageni strain FIOCRUZ L1-130 (LIC) (i/p). Graphs represent the percentage of CD4⁺ and CD8⁺ T cells and their subpopulations at 30 days post-infection. Each dot represents one animal (n=3 per group). Statistical analysis was performed using two-way ANOVA, followed by Tukey’s test, with familiar α of 0.95, *p-*values: *< 0.05; **< 0.01; \*\*\**p*< 0.001; \*\*\*\**p*< 0.0001; and ns: non-significant. Mice were obtained from the Animal Care Unit from UTHSC.

## Discussion

This study investigates the role of murine C3 during tissue colonization of pathogenic *L. interrogans* and its potential contribution to kidney fibrosis during chronic infection. Establishing a model for live, chronic leptospiral infection is a complex but critical step toward understanding the mechanisms by which leptospirosis contributes to chronic kidney disease. Here, we provide the first report of chronic infection using C57BL/6J mice and their C3 knockout counterparts, shedding light on C3’s role in the immune response to *Leptospira*.

The lack of C3 did not affect clinical scores of LIC-infected mice even at 180 d.p.i., which was similar to their WT counterparts. This resilience is likely due to compensation by other immune mechanisms. However, kidney colonization by *L. interrogans* were apparently higher in LIC-infected C3KO mice at 30 d.p.i., where 7 out of 8 mice displayed leptospiral loads exceeding 10^3^ leptospires/mg tissue, compared to minimal colonization in WT mice (only 1 in 10 mice) (**Fig 1** and **Fig 6**). These findings suggest that while the absence of C3 does not influence disease severity, it does increase kidney colonization by *L. interrogans* in C57BL/6J mice at 30 d.p.i. Recently, our group demonstrated that during the early stages of infection with *L. interrogans* serovar Kennewinki strain Pomona Fromm, leptospiral loads were significantly higher in the absence of C3 in the kidney, liver, spleen at 3 d.p.i. and in the urine at 6 d.p.i. compared to WT-infected mice [27]. These observations indicate that C3’s role in controlling bacterial load and kidney colonization is critical during the initial phase of infection.

Kidney fibrosis - a hallmark of chronic leptospirosis - was evaluated alongside the expression of fibrosis-related genes. Although larger fibrotic areas were observed in C3KO mice at 30 d.p.i., the expression of fibrosis-related gene (e.g., *FN1*, *COL1A1*, and *α-SMA*) did not differ significantly between groups. This suggests that fibrosis formation may proceed via these biochemical pathways independently of C3. These results contrast with findings from other murine models with genetic modification, such as *iNOS* KO mice [32], where FN1 and α-SMA levels were lower after LIC infection, as well as DAF-1 KO mice [33], and humanized TLR4 mice [34], which demonstrated a higher progression of fibrosis, evidenced by increased COL1A1 expression at different time points (90 d.p.i. for DAF-1 KO mice and 15 d.p.i. for humanized TLR4 mice). Our findings suggest that C3 does not significantly influence the expression of fibrosis-related genes during LIC infection, suggesting that other pathways may play a more dominant role in fibrosis development.

C3-derived fragment C3d plays a pivotal role in immune modulation. C3d-bound antigens interact with Complement Receptor 2 (CR2, also known as CD21) on B lymphocytes, promoting their activation, memory formation, and immunoglobulin class switching [35]. Vassalakis et al. [27] demonstrated that C3KO mice immunized with 3 x 10^7^ *L. interrogans* serovar Kennewiki strain Pomona Fromm or with 5 µg of recombinant leptospiral protein LigBC produced significantly lower levels of specific IgG compared to their WT counterparts. In that study, the same animals received a booster dose two weeks later, further highlighting reduced IgG production in the absence of C3.

Based on these findings, we expected that C3KO mice infected with a single dose of 10^8^ LIC would also exhibit lower levels of specific antibodies compared to WT LIC-infected mice. However, specific antibody levels remained similar or, in some cases, even higher despite the absence of C3. One possible explanation is that C3 deficiency allows leptospires to persist for a longer period, leading to increased bacterial multiplication and, consequently, a higher antigen load. Another possibility is that the higher infectious dose used in this study may have masked differences in antibody production, as such differences are typically more pronounced at lower antigen doses [36] or during secondary immune responses [37].

Interestingly, after 30 d.p.i., the IgG response was predominantly composed of IgG3. In mice, IgG3 is associated with T-cell independent responses against lipopolysaccharide (LPS) antigens [38–40], whereas in humans, this response is driven by IgG2 [41]. This suggests that the absence of C3d may influence the T-cell independent response against LPS.

Proteome profiling revealed minimal differences in cytokine levels between infected and control groups, with the exception of IL-16 - a T CD4^+^ chemoattractant [42]. Chemokine levels also remained largely unchanged, except in WT LIC-infected mice, which exhibited lower levels of M -CSF, sICAM-1, and SDF-1, and in C3KO LIC-infected mice, which showed reduced M-CSF and elevated SDF-1. These variations may reflect the cytokine and chemokine activity during leptospiral dissemination [31], which appears to normalize during chronic kidney colonization. Interestingly, C5/C5a levels were reduced, though not eliminated, in C3KO mice when compared to WT control. This supports the hypothesis that murine thrombin can directly cleave C5 into C5a and C5b fragments [43–45], enabling the formation of MAC (C5b-9_n_) despite the absence of C3.

Contrary to expectations, chronic LIC infection had only a minor impact on renal functional parameters. These findings are consistent with findings in galectin-3 knockout mice [46], where no significant deviations in renal biomarkers were observed. Such deviations may be unique to human leptospirosis [47, 48], as murine models appear to be more resilient to these changes. In leptospirosis patients, hypokalemia is commonly observed due to excessive urinary potassium excretion [49, 50]. Measuring potassium levels in murine models of leptospirosis could provide valuable insights and should be considered in future studies.

We aimed to model acute kidney injury (AKI) and compare its effects to kidney injury caused by LIC infection. Cisplatin, a chemotherapeutic agent known for its potent nephrotoxicity, frequently induces AKI in patients, leading to treatment interruptions in 30-40% of cases [51, 52]. To assess its impact in conjunction with LIC infection, we administered cisplatin to mice and analyzed the resulting kidney damage. While C3 was not required to exacerbate cisplatin-induced damage, LIC infection led to a higher inflammation score when animals were pre-treated with cisplatin. However, we did not assess whether cisplatin persists in the organism beyond one month or whether it influences LIC colonization and growth in the kidney tubules, which could potentially explain the reduced LIC load in some mice. Further experiments are needed to clarify these findings.

The cellular immune response to pathogenic *Leptospira* also remains poorly understood. Here, we analyzed the lymphocyte populations after 30 d.p.i.. In the absence of C3, *L. interrogans* infection led to a relatively discrete percent reduction in number of cytotoxic T CD8^+^ lymphocytes in the kidney and lymph nodes but not in the spleen, which may contribute to chronic local *Leptospira* infection. Pathogenic *L. interrogans* is known to induce cell death processes such as necroptosis in different immune cells [53], a process that exacerbates persistent inflammation during the early stages of infection. Blocking the necroptosis pathway has been shown to attenuate the effects of acute leptospirosis [54]. This necroptosis mechanism may account for the lower abundance of effector cytotoxic T CD8^+^ lymphocytes during chronic infection, allowing leptospires to persist in the kidney, which serves as the primary site of inflammation during leptospirosis. Further studies are needed to fully elucidate this phenomenon.

Additionally, LIC-infected C3KO mice exhibited a higher proportion of naïve T lymphocytes and a lower proportion of early effector CD4^+^ and CD8^+^ T lymphocytes in the kidney and peripheral lymph nodes. C3 is produced locally during interactions between antigen-presenting cells and T lymphocytes, where it provides costimulatory signals and maintains the viability of naïve T cells [55–57]. It also supports the activation and expansion of CD4^+^ and CD8^+^ T lymphocytes in response to viral [58, 59] and bacterial [60] infections. Moreover, the local production of C3 appears critical for differentiation of naïve T lymphocytes into effector cells; in its absence, C3KO mice retain a relatively elevated proportion of naïve T cells, underscoring the importance of C3 in promoting the differentiation and expansion of effector T cell populations necessary to combat infection.

Overall, the absence of C3 does not impact the mouse survival during leptospirosis. However, one-month post-infection appears to be a critical time point in murine leptospirosis if C3 deficiency is present, as both leptospiral load in the kidney and fibrosis formation were more pronounced at this stage. Additionally, murine C3 deficiency impairs the differentiation of naïve T cells into effector helper and cytotoxic T lymphocytes in the kidney and lymph nodes after 30 d.p.i., suggesting that the depletion of cytotoxic T lymphocytes in the absence of C3 may contribute to chronic leptospirosis. Our findings highlight the potential role of C3 in regulating effector helper and cytotoxic T lymphocytes, serving as a key link between the innate and adaptive immune system in leptospirosis.

## Materials and Methods

### Animals

Male, 10-week-old C57BL/6J wild type (WT) (RRID:IMSR_JAX:000664) and B6.129S4-C3^tm1Crr^/J (RRID:IMSR_JAX:029661) congenic homozygous C3-deficient (C3KO) mice were purchased from The Jackson Laboratory (Bar Harbor, ME). Animals were maintained and used in a specific pathogen-free environment at the Animal Care Facilities at the Institute of Biomedical Sciences of the University of São Paulo (ICB – USP) or at the University of Tennessee Health Science Center (UTHSC), as indicated in each figure legend.

### Ethics statement

All animal procedures carried out in this study were approved by the Ethical Committees of ICB-USP (Committee Protocol number: 9917191218) and UTHSC (Committee Protocol number: 22-0362).

### Leptospira infection

Pathogenic *L. interrogans* serovar Copenhageni strain FIOCRUZ L1-130 (LIC) were acquired from the Bacterial Zoonosis Laboratory at the Medicine Veterinary School at University of São Paulo or from the UTHSC. LIC was propagated after passage in hamster to maintain virulence. LIC was thawed from −80°C storage and cultured in Hornsby-Alt-Nally (HAN) semi-solid medium [28] at 29°C. For subsequent passages, 1 ml of the HAN culture was transferred into 9 ml of Difco *Leptospira* Medium Base EMJH (Ellinghausen–McCullough–Johnson–Harris, BD, ref. 279410) supplemented with Difco Enrichment EMJH (BD, ref. 279510). To preserve virulence, cultures were limited to two passages. Bacterial cells were harvested by centrifugation at 5660 *x g* for 20 min at 20°C, resuspended in 10 ml sterile PBS, and washed twice under the same conditions. Leptospires were quantified using a Petroff-Hausser chamber under a dark-field microscope (ZEISS Scope. A1). A total of 10^8^ leptospires in 150 μl sterile endotoxin-free Dulbecco’s PBS (LOT # 3303187, Millipore Corp, Billerica MA, USA) was injected intraperitoneally (i/p). Control mice received an equivalent volume of PBS and kept under the same experimental conditions.

### Acute kidney injury with cisplatin injection

WT and C3KO mice received a single injection (i/p) of cisplatin (10mg/kg) (Millipore-SIGMA, #PHR1624) diluted in 150 μL sterile PBS. Control groups received 150 μL PBS alone. After one month, mice were further injected with 150 μL PBS (control) or with 10^8^ LIC in 150 μL (i/p). Animals were monitored daily for weight changes and clinical signs for 15 d.p.i. and then euthanized with isoflurane. Kidney, liver and lung samples were collected for histopathological analysis; and part of them were treated with RNAprotect tissue reagent (QIAGEN, ID: 76106) before DNA extraction.

### Histopathological analysis

Kidney and liver samples were immediately placed in a 4% formaldehyde and later transferred to 70% ethanol before processing. Tissue sections were fixed with Dubosq-Brasil solution (1 g picric acid; 150 mL of 80% ethanol; 60 mL of 40% formalin; and 15 mL of glacial acetic acid) for 2 h and post-fixed with 10% formalin (aqueous formaldehyde), and embedded in paraffin. Sections were stained with hematoxylin and eosin (HE) or immersed for 30 min in a 0.1% Sirius Red solution, followed by hematoxylin staining. Sirius Red staining highlights the presence of collagen type I and III fibers. An inflammatory score was assigned to the kidney and liver based on the following criteria: 0 – No irregularities; 1 – Infiltrates detected, presence of apoptotic cells < 25% of the tissue. For kidney samples, nephritis and visible fibrosis were also noted; 2 – Infiltrates detected, with apoptotic cells in > 25% but < 50% of the tissue. For kidney samples, nephritis and visible fibrosis were observed. For liver samples, regeneration marks, duplicated nuclei and hepatocyte destrabeculation were identified; 3 – Infiltrates detected, with apoptotic cells in > 25% but < 50% of the tissue. For kidney samples, nephritis and visible fibrosis were present. For liver samples, regeneration marks, duplicated nuclei and hepatocyte destrabeculation were identified.

### Fibrosis quantification

Sirius Red-stained kidney sections were scanned with the ZEISS Axioscan 7 slide scanner. Images were analyzed in ZEISS ZEN BLUE 3.1 software. Red intensity was quantified using Image J software, with the green channel of the RGB stack command and a threshold of 0-110.

### q-PCR and RT-PCR

The presence of LIC DNA in the liver and kidney was quantified by qPCR using a TAMRA probe (sequence: 5’-CTCACCAAGGCGACGATCGGTAGC-3’, FAM-TAMRA; Eurofins) and primers (sequences: F 5’-CCCGCGTCCGATTAG-3’ and R 5’-TCCATTGTGGCCGAACAC-3’; Eurofins) targeting leptospiral *16S rRNA*. DNA was extracted using the NucleoSpin® Tissue kit (#740952.250, Clontech, Mountain View, CA), according to the manufacturer’s instructions. DNA concentration and purity were measured at 260/280 nm and 260/230 nm using a NanoDrop instrument (Thermo Scientific). qPCR was conducted alongside a standard curve of 10^6^ to 1 *L. interrogans* DNA copies.

Total cellular mRNA from the kidney tissue was extracted using RNeasy® Mini Kit (#74104, Qiagen, Germany). RNA concentration and purity were assessed at 260/280 nm before proceeding with cDNA production. A high-capacity cDNA reverse transcriptase kit (#4368814, Applied Biosystems) was used for cDNA preparation. TAMRA probes and primers for *COL1A1*, (sequences: F 5’-TAAGGGTACCGCTGGAGAAC-3’, R 5’-GTTCACCTCTCTCACCAGCA-3’ and TAMRA probe 5’-AGAGCGAGGCCTTCCCGGAC-3’, FAM-TAMRA; Eurofins) *iNOS* (sequences: F 5’-GCTGGGCTGTACAAACCTTC-3’, R 5’-GCATTGGAAGTAGAAGCGTTTC −3’ and TAMRA probe 5’-GGCAGCCTGTGAGACCTTTGAT-3’, FAM-TAMRA; Eurofins) and *β-actin* (endogenous control, sequences: F 5’-CCACAGCTGAGAGGGAAATC-3’, R 5’-CCAATAGTGATGACCTGGCCG-3’ and TAMRA probe 5’-GGAGATGGCCACTGCCGCATC-3’, FAM-TAMRA; Eurofins) were used, along with Taqman Gene Expression Assays kits for *FN1* and *ACTA1* (Mm01546133_m1 and Mm01256744_m1, Thermo Fisher Scientific).

The qPCR mixture contained a final mixture of 900nM for each primer, 250nM for the specific TAMRA probe, 10 μl of TaqMan^TM^ Fast Advanced Master Mix (#4444557, Applied Biosystems) and 2 μl of the DNA or cDNA sample, for a volume of 20 μl. All reactions were performed in duplicate. The amplification was done in a QuantStudio 3 equipment, following the protocol: 2 min at 50°C, 20 s at 95°C, followed by 40 cycles of amplification (1 s at 95°C and 20 s at 60°C). Data for gene expression was analyzed with ΔΔCt comparative method.

### Flow cytometry

Spleen, kidney and six peripheral lymph nodes were collected from three mice per group. Cell suspensions were prepared, treated with red blood cells lysis solution using a previously described protocol with some modifications [29]. Dead cells were excluded using a live/dead cell stain and counted with a Luna Cell Counter (Logos Biosystems, South Korea). Approximately 10^6^ cells were seeded per well in a 96-well microtiter plate and blocked with anti-mouse CD16/32 antibody (1:100) for 15-20 min on ice in PBS, pH 7.5. Cells were washed with cell staining buffer [Ca^2+^-and Mg^2+^-free PBS containing 3% heat-inactivated fetal bovine serum, 0.09% sodium azide (Sigma-Aldrich, St. Louis, MO), 5 mM EDTA]. Surface staining was performed with primary conjugated antibodies against various cell surface markers, incubated for 30 min at 4°C in the dark, and washed twice with cell staining buffer. Cells were fixed with 4% paraformaldehyde for 10 min followed by a single PBS wash. Fixed cells were finally resuspended in cell staining buffer and acquisition was performed using a BioRad ZE5 Cell analyzer. The data was analyzed using FlowJo software.

### Proteome Profile Array

Mouse serum cytokines, including some chemokines, and Complement C5/C5a were detected using the Proteome Profiler Array Cytokine Mouse Kit (Panel A, catalog ARY006, R&D Systems, Minneapolis, MN, USA) [30, 31] following the manufacturer’s instructions. Array membranes were first treated with blocking buffer (provided by the manufacturer) for 1 h in room temperature in a rotary shaker. Approximately 100 μl pooled serum per group were mixed for 1 h with a cocktail of 40 different biotinylated detection antibodies and then incubated overnight at 4°C with the Mouse Cytokine Array nitrocellulose membrane. The membranes contained spotted capture antibodies in duplicate and washed to remove any non-binding conjugates. They were then incubated with a horseradish protease-conjugated anti-mouse IgG secondary antibody. Finally, the membranes were washed three times with wash buffer and developed using the developing solution provided in the kit. Chemiluminescence intensity was proportional to the amount of cytokine bound to the membrane. Images at different exposure times were captured using a Chemi-Doc image analyzer, and mean pixel density was quantified using QuickSpots software.

### Serum analysis of creatinine, uric acid and urea

After euthanasia, mouse blood was collected through heart puncture and set to rest for 1 h in ice. The blood was centrifuged at 1500 x *g*, and the serum was harvested and stored at −80°C until later use. Serum creatinine (# 27-500), uric acid (# 140-1/100) and urea (#: 96-300) were analyzed with Labtest kits (Minas Gerais, Brazil). Colorimetric tests were performed with 10 µl serum and compared to a standard solution (urea – 70 mg/dl; uric acid – 4mg/dl; and creatinine – 4mg/dl) provided by the manufacturer. Absorbance was measured using a Ultrospec 2100 Pro equipment (GE Healthcare, Chicago, USA). Urea, uric acid and creatinine levels were determined at 600 nm, 505 nm, and 510 nm, respectively.

### Serum immunoglobulin analysis

The concentrations of serum specific anti-LIC IgM, IgG and IgG subtypes (IgG1, IgG2b and IgG3) were measured by ELISA. We utilized 10^6^ heat-killed LIC in 100 μl of carbonate buffer (Carbonate Coating Buffer CB01100, Thermo Scientific^TM^) to coat 96-wells plates (MaxiSorp, Thermo Fisher Scientific, Waltham, MA) overnight at 4°C. The plates were washed 4 times with 300 μL of PBS + 0.05% Tween (PBS-T) and incubated with 250 μL of blocking buffer (PBS-T + 1% fresh bovine serum albumin) for 1h at 37°C.

Followed by washing with PBS-T, the plates were dried and incubated for 1h at 37°C with mouse serum (1/100) diluted in 100 μL of blocking buffer. After another washing with PBS-T, these plates were incubated with rabbit secondary horseradish peroxidase-conjugated (HRP) antibodies, specific against mice IgM, IgG, IgG1, IgG2b or IgG3 (1/10000, Jackson ImmunoResearch, PA). 3,3’,5,5’-tetramethylbenzidine (TMB, substract solution N301, Thermo Scientific^TM^) and a stopping solution (N600, Thermo Scientific^TM^) were added to the wells and optic density was determined with the SpectraMax Plus 384 ELISA reader spectrophotometer at 450 nM wavelength.

### Statistical analysis

Statistical analyses were performed using GraphPad Prism software or JASP (v. 0.19.1), and graphs were prepared with GraphPad on Windows 11. Prior to conducting ANOVA tests, groups were evaluated for normal population distribution using a Shapiro-Wilk test. Two-way ANOVA was performed, considering two independent factors: mouse genotype (WT or C3KO) and treatment type (control or LIC). When appropriate, post-hoc multiple comparisons were conducted using Tukey’s correction were applied, with a family-wise alpha of 0.05. If ANOVA was not applicable, the Kruskal-Wallis nonparametric test was used, followed by Dunn’s post-hoc tests.

## Acknowledgments

We express our sincere gratitude to Dr. Gláucia Maria Machado-Santelli (ICB-USP), Dr. Diedre Daria and Dr. Tony Marion of the Flow Cytometry and Cell Sorting Core facility at UTHSC. We would like to thank Dr. Angela Silva Barbosa (Institute Butantan, São Paulo) for technical support. We also would like to acknowledge Dr. Ae-Kyung Yi’s lab for access to the Chemi-Doc system for imaging Western blots. This work was supported by Fundação de Amparo à Pesquisa do Estado de São Paulo (FAPESP) Proc.# 2017/12924-3, Proc. #2018/26574-7, and Proc.# 2022/05793-8, by Conselho Nacional de Pesquisa e Desenvolvimento Tecnológico (CNPq) Proc.# 312654/2021-9, and by the United States National Institute of Allergy and Infectious Diseases (NIAID), National Institutes of Health (NIH), grant numbers R01AI139267 and R44AI167605 to MGS.

---

**S1 Fig.**
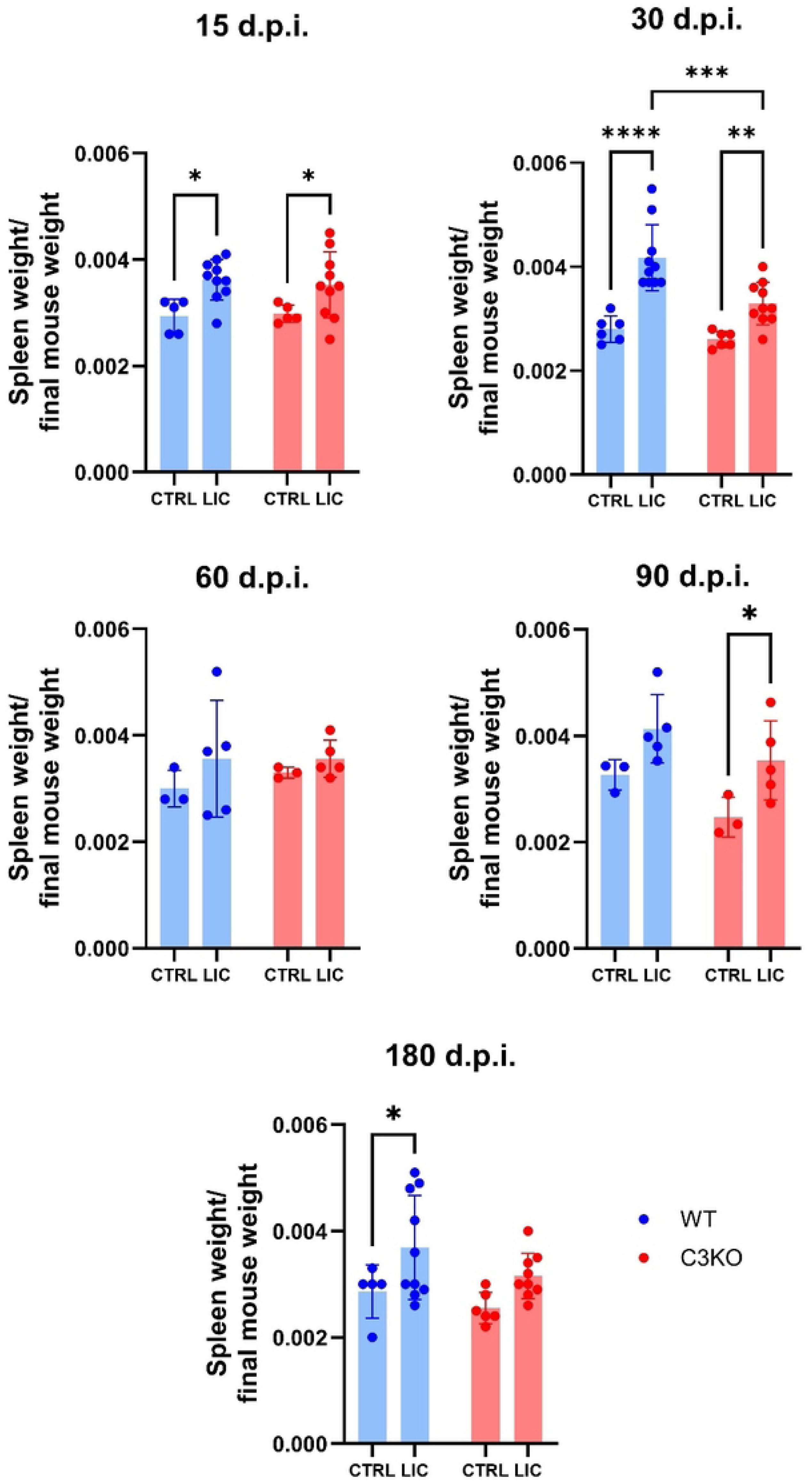
Spleen-to-total body weight ratio. The spleen weight of WT and C3KO mice, injected with PBS (control) or 10^8^ *L interrogans* serovar Copenhageni strain FIOCRUZ L1-130 (LIC) (i/p) was measured after euthanasia and compared to the respective total body weight. Each dot represents one animal. Before statistical analysis, variance homogeneity of and normality of the groups was verified. A two-way ANOVA was performed, followed by Tukey’s post-hoc test. Significance was set at α = 0.05. \**p* < 0,05; \*\**p* < 0,01; \*\*\**p* < 0,005, and \*\*\*\**p* < 0,001. Mice were obtained from the Animal Care Unit from ICB-USP. Infections at 15, 30, 90 and 180 d.p.i. were repeated twice, while infection at 60 d.p.i. was performed once.

**S2 Fig.**
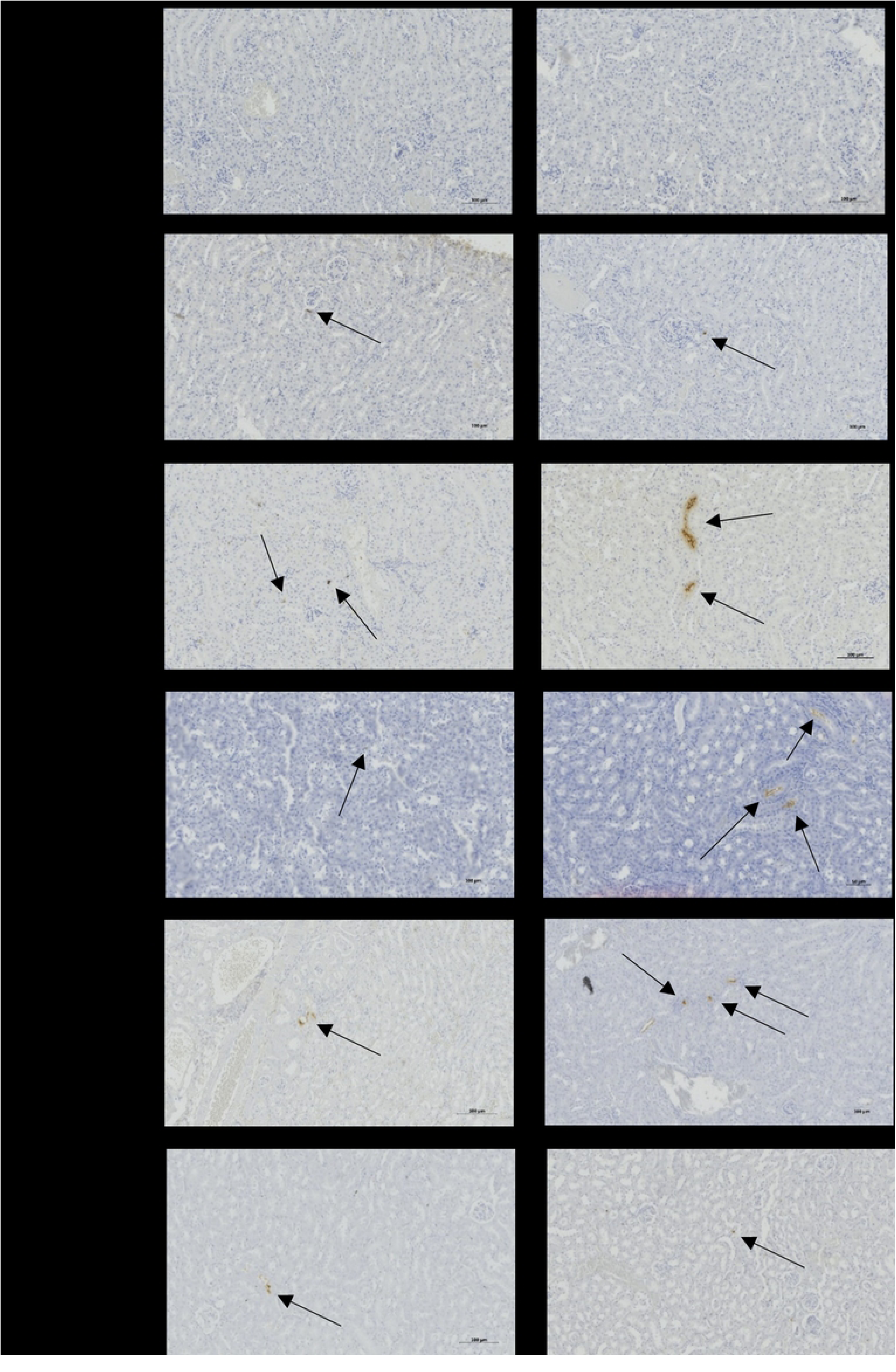
Immunohistochemistry analysis after LIC infection. WT and C3KO mice were inoculated with PBS (CTRL) or 10^8^ *L interrogans* serovar Copenhageni strain FIOCRUZ L1-130 (LIC) (i/p). The presence of LIC antigen (stained brown and indicated by arrows) was detected in all kidneys from LIC infected mice after incubation with rabbit anti-*Leptospira* polyclonal antibodies. Images were captured at 200 x magnification. Mice were obtained from the Animal Care Unit from ICB-USP.

**S3 Fig.**
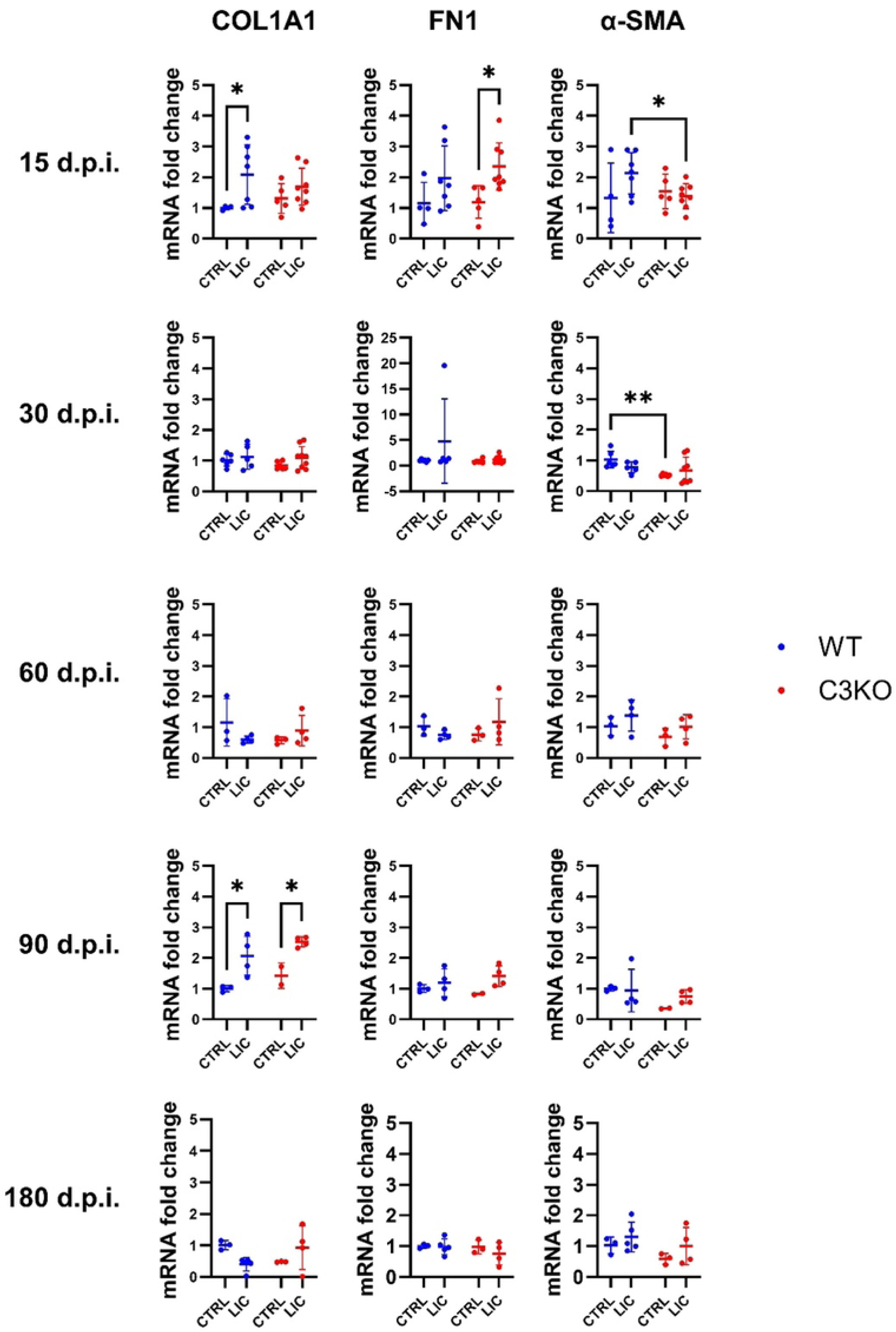
Gene expression of fibrosis-related genes. WT and C3KO mice were inoculated with PBS (control; CTRL) or 10^8^ *L interrogans* serovar Copenhageni strain FIOCRUZ L1-130 (LIC) (i/p). After 15, 30, 60, 90 and 180 days, total RNA was extracted from kidney samples to build cDNA from mRNA. We quantified the expression of COL1A1, FN1 and α-SMA through ΔΔCt comparative method, keeping β-actin as the endogenous control. A two-way ANOVA was performed, followed by Tukey’s post-hoc test. Significance was set at α = 0.05. \**p* < 0,05; \*\**p* < 0,01. Mice were obtained from the Animal Care Unit from ICB-USP.

**S4 Fig.**
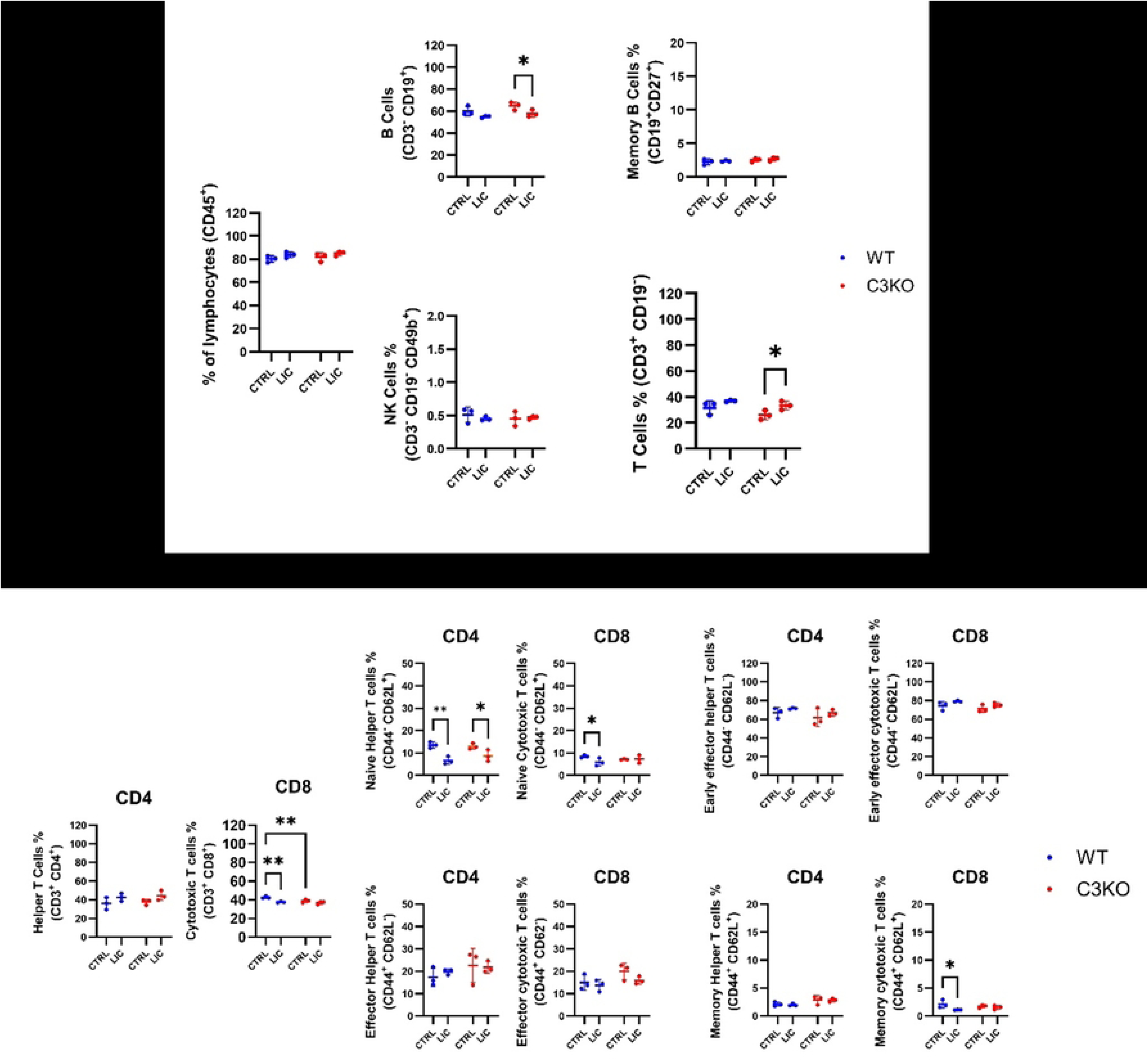
Immune cell populations in the spleen. WT or C3KO mice were inoculated with PBS (control; CTRL) or with 10^8^ *L interrogans* serovar Copenhageni strain FIOCRUZ L1-130 (LIC) (i/p). Graphs represent the percentage of CD45^+^ T and B lymphocytes, their subpopulations, and NK cells after 30 days post-infection. Each dot represents one animal (n =3 per group). Statistical analysis was performed using two-way ANOVA, followed by Tukey’s test, with familiar α of 0.95. *p-*values: *< 0.05; **< 0.01; ns = non-significant. Mice were obtained from the Animal Care Unit from UTHSC.

